# The Hidden Disorder Divide: Reconciling Benchmark Inconsistencies in Intrinsically Disordered Protein Binding Site Prediction

**DOI:** 10.64898/2026.06.24.733783

**Authors:** Nawar Malhis, Mahta Mehdiabadi, Gábor Erdős, Jörg Gsponer, Lukasz Kurgan, Silvio C E Tosatto, Zsuzsanna Dosztanyi, Damiano Piovesan

## Abstract

Computational predictors of protein-binding sites within intrinsically disordered regions (IDRs) show highly inconsistent performance across high-quality benchmark datasets. To understand the origins of these discrepancies, we systematically compared predictors across three independent test sets: two CAID datasets updated with the latest DisProt annotations and a composite dataset (DBs) assembled from DIBS, FuzDB, IDEAL, and MFIB. Predictors trained predominantly on DisProt data achieved substantially higher AUCs on the CAID sets but performed poorly on the DBs. In contrast, predictors trained on older, low-quality PDB-based datasets showed balanced performance across all sets, with a slight preference for DBs. Predictors with mixed training exposure displayed intermediate behavior. Through controlled experiments using identical CNN architectures and feature analysis, we demonstrate that the dominant factor driving these performance differences is the intrinsic disorder propensity of the binding sites themselves. Binding residues in DisProt-based datasets exhibit markedly higher average disorder propensity scores than those in PDB-derived datasets. This previously unrecognized selection bias — literature studies preferentially characterizing more disordered binding sites, while PDB-derived annotations capture less disordered ones — effectively splits IDR-protein binding sites into two distinct categories. Predictors optimized on one category therefore generalize poorly to the other. Binding-site length and sequence conservation play only minor or negligible roles in explaining the observed inconsistencies. These findings highlight a critical limitation in current benchmarking practices and training strategies for IDR-binding site prediction, underscoring the need for more balanced and disorder-aware reference datasets. Finally, the diagnostic techniques introduced here could prove valuable beyond the specific application examined in this study.

The data and code used to generate all figures and tables are available at: https://github.com/NawarMalhis/HDD

## 1. Introduction

Intrinsically disordered regions (IDRs) in proteins are segments that lack a stable three-dimensional structure under physiological conditions. These regions are widespread, and proteins with extensive IDRs account for approximately one-third of eukaryotic proteomes [1]. IDRs play critical roles in diverse cellular interactions, supporting essential biological processes [2, 3, 4]. Unlike folded protein domains, whose interactions depend on complementary shapes and limited flexibility, the structural plasticity of IDRs enables them to adapt to a wide range of binding partners [5, 6]. Exposure of IDRs to alternative splicing [7] and post-translational modifications further contributes to the regulation and diversification of protein interactions.

In addition, IDRs often harbor multiple binding sites within the same region, enhancing their versatility. Their interactions are typically specific yet of low affinity [8], making them reversible and particularly suited for regulatory roles in signaling pathways [3]. Accordingly, IDRs are enriched in eukaryotic hub proteins [9].

IDRs interact with partners through diverse mechanisms mediated by specialized elements. Subclasses include Eukaryotic Linear Motifs (ELMs) and Short Linear Motifs (SLiMs) [10, 11, 12], which are short, conserved sequences (typically 3–11 amino acids) that recognize specific domains in partner proteins. Molecular recognition features (MoRFs) [13, 14, 15] consist of disordered segments up to 70 residues and also transition to ordered structures upon binding. Despite their diversity, most of these sites share common traits, such as some degree of sequence conservation. Advances in computational biology have fueled efforts to predict IDR protein-binding site locations directly from sequence data, leading to a broad range of prediction methods [15–50]. These tools have been instrumental in identifying potential interaction sites and deepening our understanding of IDR functionality [51–57].

The Critical Assessment of Protein Intrinsic Disorder (CAID) is a community-wide initiative that benchmarks computational methods for predicting IDRs and their binding sites. Three assessments were held in 2018 [58], 2022 [59], and 2024 [60], each using the DisProt database [61] as a reference. Nevertheless, several tools show uneven performance across the three CAID rounds, with even greater inconsistencies emerging when tested against other high-quality, manually curated databases. Notably, these inconsistencies were observed only for binding, not for general disorder. In this study, we examine how dataset composition and predictor characteristics influence the observed performance of IDR–protein-binding predictors.

### 2. Datasets and existing predictors

CAID employs a blind-test approach in which methods are evaluated on proteins for which DisProt annotations are only available after the challenge submission deadline. This design minimizes the risk that target proteins appear in the training sets of methods optimized on DisProt data, thereby reducing potential contamination. Because CAID2 and CAID3 contain few IDR-protein-binding annotations, we merged them into a single dataset, denoted CAID2&3. We filtered sequences in CAID1 and CAID2&3 that share more than 30% sequence identity. We updated these datasets using IDR–protein binding site annotations from the DisProt 2025_06 release and appended a “u” suffix to denote the updated versions, i.e., CAID1u and CAID2&3u. Additionally, we assembled a third test dataset (DBs) from four high-quality, manually curated sources: DIBS [62], FuzDB [63], IDEAL [64], and MFIB [65]. DBs sequences were filtered to ensure <30% identity with both the TR2008 training dataset (used to train several predictors) and the DisProt 2025_06 database. Our study, therefore, relies on three independent test sets: CAID1u, CAID2&3u, and DBs. While all annotations in DBs derive from PDB structures, those in DisProt (including the CAID datasets) are primarily literature-based and include binding evidence from alternative experimental types. Further details on the datasets are provided in **Supplementary Note 1** and **Supplementary Table 1**.

Using these three test sets, we evaluated MoRFchibi, fMoRFpred, and the nine additional IDR–protein-binding predictors from CAID2 that have associated publications or preprints. For fMoRFpred, we generated predictions using its HTML server, and we used the CAID Portal server to generate predictions for the remaining predictors. **Table 1** summarizes their AUC performance, focusing on within-predictor comparisons across the datasets. Predictors that did not participate in the CAID1 round were evaluated only on the CAID2&3u and DBs datasets.

We divided the 11 predictors into three groups based on their training datasets. Group A includes the five tools trained on DisProt datasets: DeepDRPBind-protein, DisoRDPbind-protein, DeepDISObind-protein, DRPBind-protein, and AlphaFold-binding. These exhibit markedly higher performance on the CAID datasets than on DBs. Group B comprises fMoRFpred, MoRFchibi, and OPAL, all of which are trained on the TR2008 dataset derived from PDB structures. These predictors perform comparably on the CAID and DBs datasets, with a slight (often trivial) advantage on DBs. Group C consists of MoRFchibi-light and MoRFchibi-web. Although trained on TR2008, both incorporate the ESpritz predictor and are thus directly exposed to DisProt data; consequently, they perform better on the CAID datasets than on DBs, but with less pronounced contrast than Group A. Finally, ANCHOR-2 [35] is not assigned to any group. Despite being trained on PDB-derived data similar to Group B, its performance trends align more closely with those of Group A.

Interestingly, the weak performance of Group A predictors on the DBs dataset is accompanied by abnormal ROC curves for most Group A predictors, suggesting inconsistencies between their DisProt-based training data and the DBs datasets. In contrast, the shapes of the ROC curves for Group B predictors against CAID2&3u appear unaffected. ANCHOR-2 shows ROC curve patterns similar to Group A. (**Figure 1**; **Supplementary Figure 1**)

**Table 1.** The AUC values in identifying IDR-protein binding sites for all 11 predictors against our three test datasets. Tools constructed after the CAID round 1 event are not evaluated on the CAID1u dataset.

| Predictor | Group | AUC | | | $\Delta$ AUC | | |
| --- | --- | --- | --- | --- | --- | --- | --- |
|  |  | CAID1u | CAID2&3u | DBs | CAID2&3u - CAID1u | DBs - CAID1u | DBs - CAID2&3u |
| AlphaFold-binding | A |  | 0.787 | 0.412 |  |  | -0.376 |
| ANCHOR-2 |  | 0.739 | 0.755 | 0.388 | 0.016 | -0.350 | -0.367 |
| DeepDISObind-protein | A |  | 0.760 | 0.393 |  |  | -0.366 |
| DeepDRPBind-protein | A |  | 0.719 | 0.577 |  |  | -0.143 |
| DisoRDPbind-protein | A | 0.741 | 0.765 | 0.370 | 0.024 | -0.371 | -0.395 |
| DRPBind-protein | A |  | 0.742 | 0.344 |  |  | -0.398 |
| fMoRFPred | B | 0.545 | 0.557 | 0.547 | 0.012 | 0.002 | -0.010 |
| OPAL | B | 0.665 | 0.712 | 0.733 | 0.047 | 0.067 | 0.020 |
| MoRFchibi | B | 0.544 | 0.563 | 0.612 | 0.018 | 0.068 | 0.050 |
| MoRFchibi-light | C | 0.711 | 0.774 | 0.607 | 0.063 | -0.104 | -0.167 |
| MoRFchibi-web | C | 0.685 | 0.763 | 0.635 | 0.079 | -0.049 | -0.128 |

**Figure 1.**
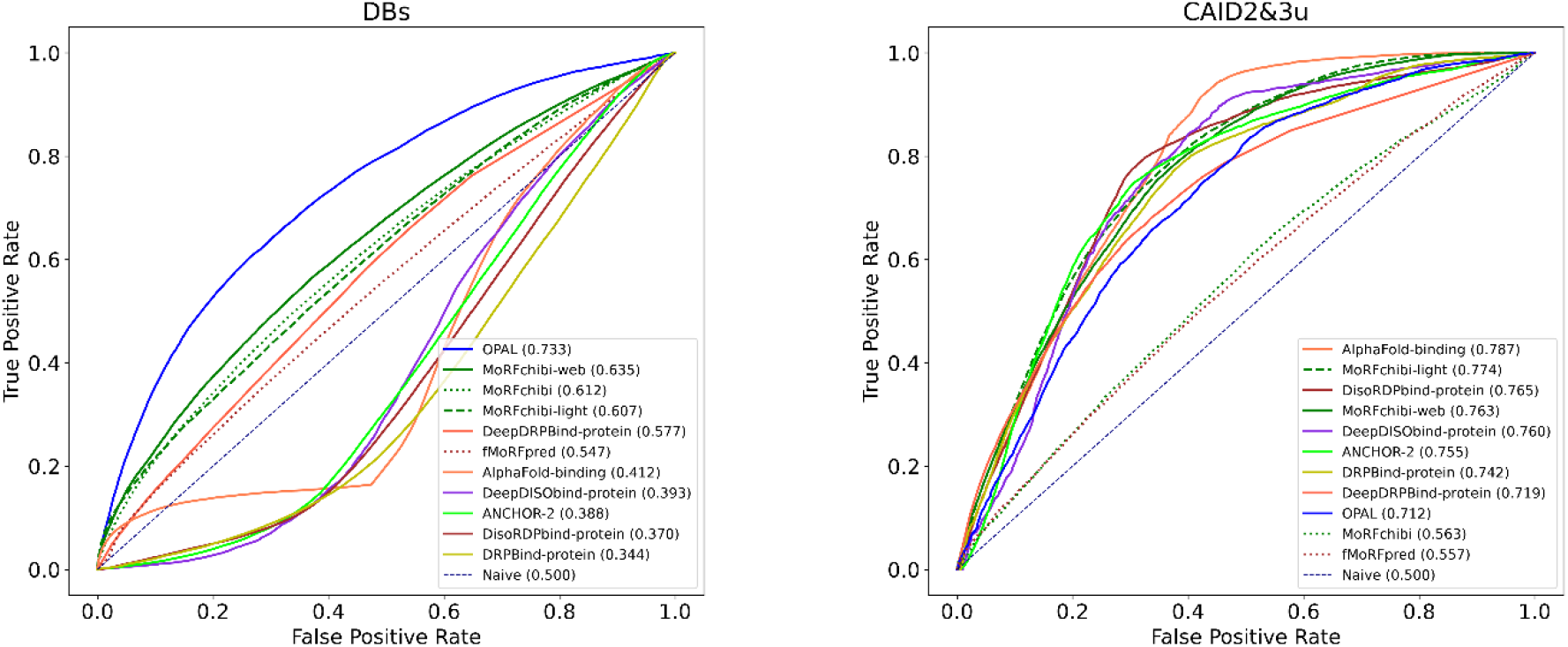
The ROC curves for the eleven IDR-protein binding predictors against the DBs (left) and the CAID2&3 (right) datasets.

### 3. Control predictors

As noted in the previous section, proteins in the CAID datasets received DisProt annotations only after the challenge submission deadline, thereby reducing the risk of contamination for tools optimized against the DisProt database. This safeguard, however, does not apply to all predictors: fMoRFpred, OPAL, and the MoRFchibi suite were trained on the TR2008 dataset, which contains homologs of CAID sequences. To address such overlaps and to better identify the sources of performance differences across datasets, it is essential to use homology-free training and test sets while also controlling for variations in machine learning architectures and input features.

To this end, we constructed a convolutional neural network comprising three convolutional and two fully connected layers, based on the IPA predictors [50] and utilizing the same input features. Unlike IPA, which employs an ensemble of four models for each predictor, our approach uses a single model (**Supplementary Note 2**).

Using CD-HIT, we filtered the TR2008 dataset to remove sequences sharing more than 30% identity with those in CAID1u and CAID2&3u, and renamed the resulting set TR2008u. We also constructed a validation dataset (VDS) from two sources, V1 and V2. V1 sequences from DisProt 7, reannotated with protein-binding sites from the DisProt 2025_06 release, after excluding entries lacking protein-binding annotations and those sharing more than 30% identity with our three test datasets (CAID1u, CAID2&3u, and DBs). And V2 sequences from TS2008 and TS2012 (**Supplementary Note 1**) that share less than 30% identity with our three test datasets and with TR2008u.

We trained four predictors: CNN_C1u against the CAID1u, CNN_C23u against the CAID2&3u, CNN_DBs against the DBs, and CNN_TR08u against the TR2008u. CNN_C1u and CNN_C23u training data are derived from DisProt; therefore, we assigned them to Group A. CNN_DBs and CNN_TR08u are trained against datasets derived from PDB structures; thus, we assigned them to Group B.

We used the HAC software (Homology Annotation Conflict) [66] to detect local homologies within individual annotated datasets, and we applied it to the four training datasets (CAID1u, CAID2&3u, DBs, and TR2008u) used to train our CNN models. TR2008u contains a high proportion of residues with annotation conflicts or redundancies. For instance, 679 class ‘1’ residues in TR2008u are homologous to at least one class’ 0’ residue, representing 17.07% of all class ‘1’ residues. In addition, 1,615 class ‘1’ residues are homologous to at least one other class ‘1’ residue, accounting for 40.60% of the total. The remaining three datasets, CAID1u, CAID2&3u, and DBs, have near-zero annotation conflict.

To further exclude the possibility of homology contamination between our training and test datasets, we applied HAM [66] (Homology-based Annotation Masking), which masks local homologous regions across datasets. HAM was applied to each test set (CAID1u, CAID2&3u, and DBs) relative to TR2008u and the other two test sets. HAM-processed datasets are denoted with an “h” suffix (e.g., CAID1u becomes CAID1uh). We evaluated performance across the three test datasets, and AUC values are summarized in Supplementary Table 2. Results for the 11 predictors were consistent with those in Table 1 before HAM masking.

The training process of the CNN predictors yielded random local minima with different outcomes in each training round. Consequently, the performance differences among the CNN predictors across our three datasets may be influenced by this randomness.

To help us understand this issue, we trained 150 instances for each model, minimizing loss on the validation VDS dataset, and selected the instance with the 30^th^-highest AUC on the validation set to avoid overfitting the validation data. To assess the significance of variations in AUC values between datasets, we plotted a violin diagram of AUC values across the three test datasets for the 21 instances ranked 20^th^ to 40^th^ in AUC on the validation data, assuming they are equally likely to be selected, and highlighted the 30^th^ (the selected one), as shown in Figure 2.

**Figure 2.**
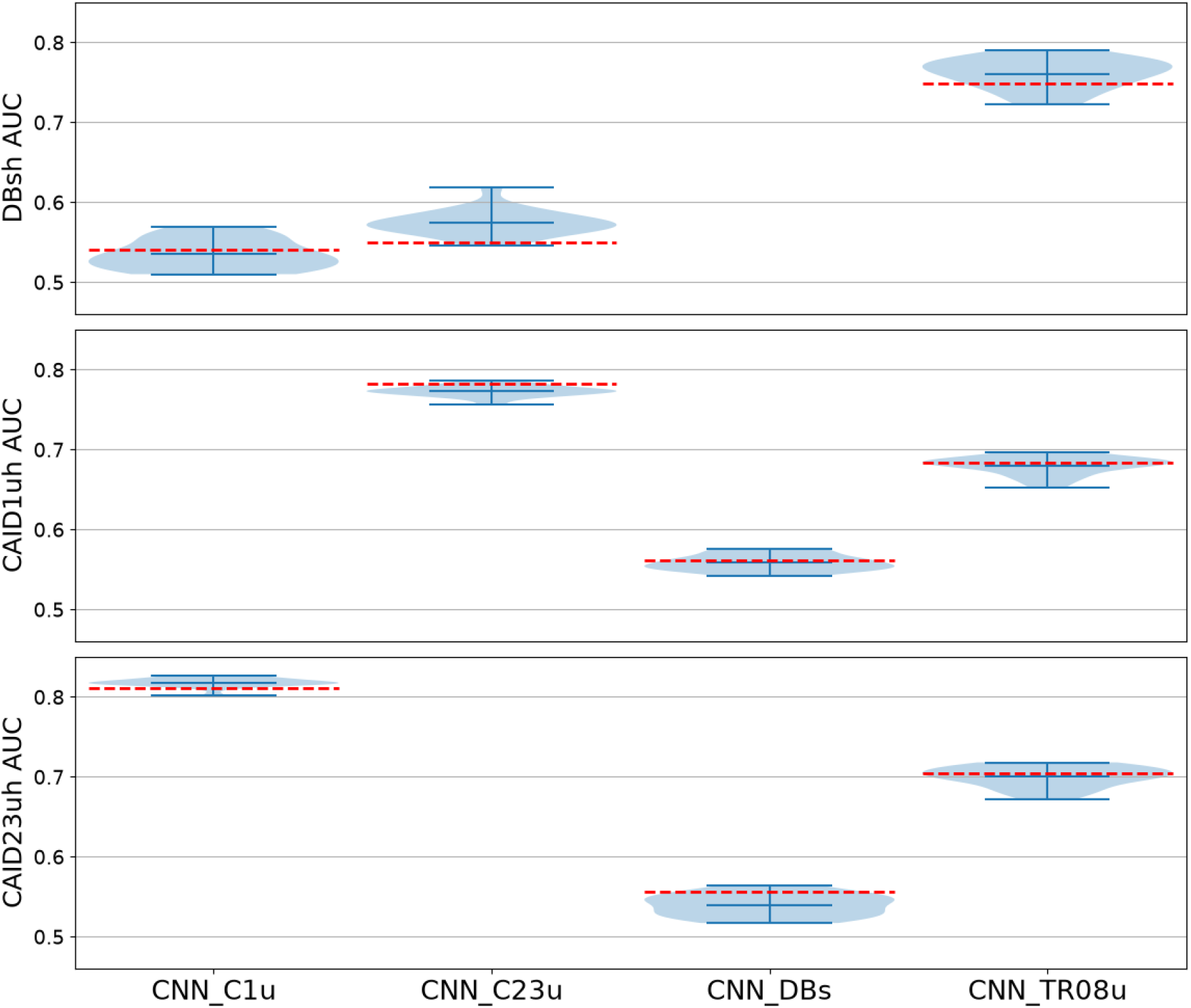
The AUC values for the selected CNN models are represented by the dotted red lines, within the distribution of the AUC values for the 21 models that are about equally likely to be chosen, represented by the shaded blue violin plots.

Figure 2. and **Supplementary Table 3** show that selecting alternative ranks—each corresponding to a different local minimum in the optimization landscape—can yield substantially different performance across datasets. For example, the selected CNN_C23u model at rank 30 (indicated by the red dashed lines in Figure 2) exhibits mixed performance compared with the other 21 models (violin plots). It lies near the lower end of the population on the DBsh dataset (AUC: 0.550) but near the upper end on the CAID1uh dataset (AUC: 0.782). In contrast, had we selected the model at rank 32, the CNN_C23u would have achieved an AUC of 0.619 on DBsh (~6 points higher) and 0.763 on CAID1uh (~2 points lower).

In essence, predictors with identical architecture, training data, input features, and hyperparameters can still exhibit markedly different AUC values on held-out test sets simply because stochastic optimization converges to different local minima. Although such stochastic instability can be mitigated through ensemble approaches (e.g., IPA), ensemble methods are not commonly used among IDR–protein binding predictors. That said, **Figure 2** illustrates that performance differences among similar models trained on different datasets are often considerably larger than these stochastic variations.

There is substantial overlap in the AUC values of CNN_DBs between the CAID1uh and CAID2&3uh datasets, as well as in the AUC values of CNN_TR08u (Figure 2). This suggests considerable similarity in the underlying characteristics of the two CAID datasets. Additionally, although both CNN_TR08u and CNN_DBs yield lower AUCs on the CAID datasets than CNN_C1u and CNN_C23u, CNN_TR08u performs markedly better across the CAID datasets than CNN_DBs. This was unexpected, given that the manually curated DBs training set is substantially larger and of higher quality than TR2008u. These findings indicate that certain IDR–protein binding features shared across specific training and test datasets strongly influence predictor performance, outweighing differences in training-set size and curation quality.

Finally, CNN_TR08u achieves a markedly higher AUC on the DBsh dataset than either CNN_C1u or CNN_C23u. This suggests that, despite clear differences between the two PDB-derived training sets (DBs and TR2008u), the disparity between the CAID datasets and the PDB-derived data is considerably larger than the differences between the PDB-derived sets themselves.

### 4. Class separation

All predictors find it easier to distinguish IDR-protein binding sites from folded domains than from other IDR regions, with which they share the property of disorder. As shown in **Supplementary Table 4**, all eleven predictors, CNN_c12u and CNN_TR08u, achieved substantially higher performance on the CAID1uh dataset when non-binding residues were restricted to those from folded domains (annotated as PDB in the CAID1 dataset) than when they were restricted to disordered residues (annotated as IDR).

In general, the AUC metric evaluates how well predicted scores separate two residue classes: IDR-protein-binding residues (class 1) and all other residues (class 0). Accordingly, training aims to assign higher scores to class 1 residues than to class 0 residues in the training data, thereby maximizing the separation between classes and improving AUC. The prediction performance on any test dataset is thus influenced by two key factors. First, separability: for example, sequence features of folded domains are more distinguishable from those of IDR-protein binding sites than are general disordered-region features. Second, dataset similarity: high AUC requires that both class 1 (binding) and class 0 (non-binding) residues in the test set resemble their counterparts in the training set. Discrepancies in the characteristics of either class reduce performance.

Consequently, to isolate the impact of class 1 differences between two datasets while neutralizing variations in class 0, we re-evaluated predictors using the class 1 residues from each dataset, paired with a shared class 0 pool from both datasets. This approach ensures that observed AUC differences between datasets primarily reflect variations in class 1. Predictors were restricted to those trained on sequences sharing <30% identity with sequences in both datasets. The AUC results in **Supplementary Tables 5(A), 6(A), and 7(A)** quantify contrasts in predictor performance on Class 1 residues after merging Class 0 residues across dataset pairs: CAID1uh and DBsh, CAID2&3uh and DBsh, and CAID1uh and CAID2&3uh, respectively. Similarly, to explicitly assess the impact of Class 0 differences, we merged Class 1 residues across datasets. The corresponding AUC results in **Supplementary Tables 5(B), 6(B), and 7(B)** quantify contrasts in predictor performance on Class 0 residues for the same pairs. **Table 2** summarizes Supplementary Table 6 by reporting the average ΔAUC for each predictor group and ANCHOR-2 across the three dataset pairs under three scenarios: standard AUC computation without any class merging; AUC after merging Class 0 residues; and AUC after merging Class 1 residues.

**Table 2.**
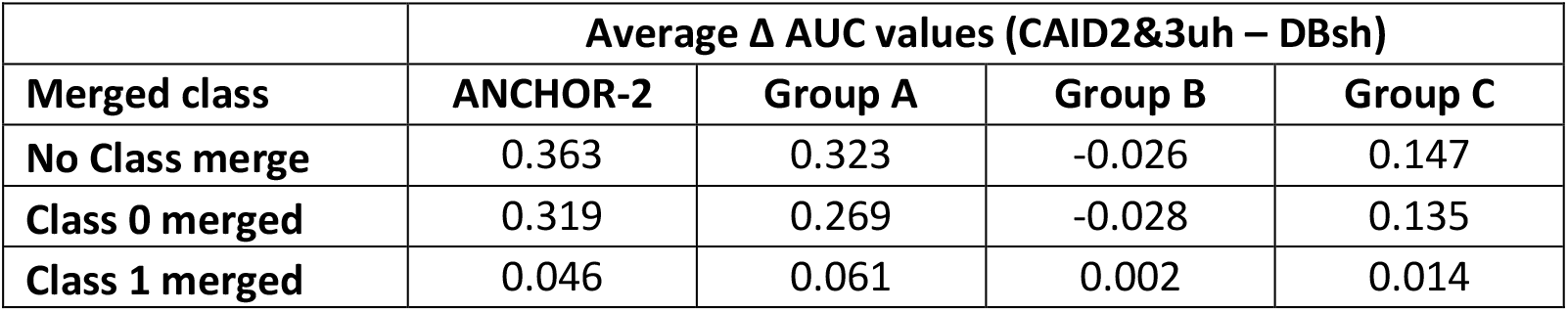
The average Δ AUC values between CAID2&3uh and DBsh for the three predictor groups and ANCHOR-2 contrasting each possible test dataset pairing in three scenarios: no class merge, class 0 merge, and class 1 merge.

| | Average $\Delta$ AUC values (CAID2&3uh – DBsh) | | | |
| --- | --- | --- | --- | --- |
| Merged class | ANCHOR-2 | Group A | Group B | Group C |
| No Class merge | 0.363 | 0.323 | -0.026 | 0.147 |
| Class 0 merged | 0.319 | 0.269 | -0.028 | 0.135 |
| Class 1 merged | 0.046 | 0.061 | 0.002 | 0.014 |

Results indicate that the substantial advantage of Group A predictors on CAID2&3uh relative to DBsh (average ~32 AUC points) stems primarily from differences in Class 1 residues, with a minor contribution from Class 0. Likewise, the small advantage of Group B predictors on DBsh relative to CAID2&3uh arises mainly from solely from Class 1 differences. Group C predictors are affected solely by Class 1 variations, while ANCHOR-2 behaves similarly to Group A. The average AUC values between CAID1uh and DBsh show a similar pattern. The ΔAUC values between CAID1uh and CAID2&3uh are limited and are likely influenced by randomness (see Section 3).

### 5. The binding site sizes

A key distinction between datasets lies in the distribution of binding residues based on binding site sizes. We use a threshold of 70 amino acids to differentiate between short and long binding sites. This threshold corresponds to the upper limit of molecular recognition features (MoRFs) and ensures that there are sufficient residues in the short category for our subsequent analysis. **Figure 3** shows that approximately 80% of the total binding residues belong to long binding sites (>70 amino acids) in the DBs dataset and the two CAID datasets, which contrasts with the TR2008u and TR2017 (used to train ANCHOR-2), where all binding sites are 30 amino acids or shorter.

**Figure 3.**
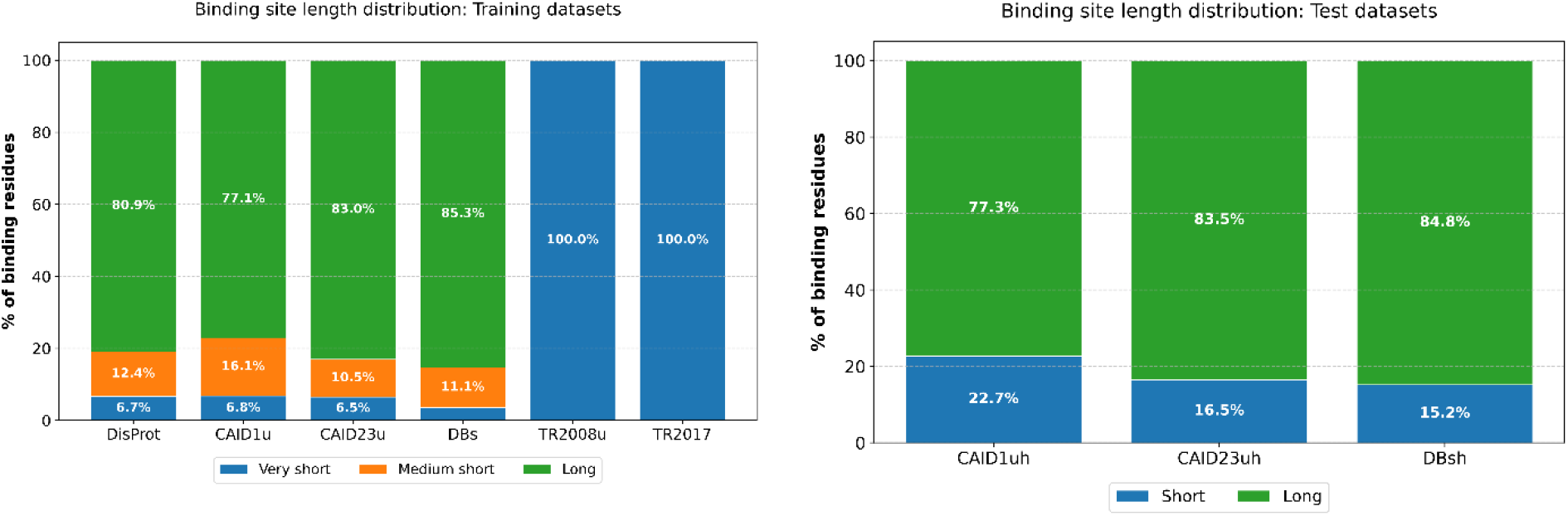
We use a threshold of 70 amino acids to differentiate between short and long binding sites. However, for the training datasets (left), we divided the short category into ‘very short’ (<= 30 amino acids) and medium short (>30 and <= 70), so that it is clear that most binding sites in the DBs, CAID1u, and CAID23u datasets are much longer than those in the TR2008u and TR2017.

**Table 3** presents AUC values for the 11 predictors and the four CNN models across the three datasets, categorizing binding sites as short (<=70 AA) or long (>70 AA) via site masking. If binding-site size significantly influences performance, predictors trained on datasets dominated by long sites would be expected to favor long sites over short ones, and vice versa. However, numerous counterexamples are observed. For instance, all Group A predictors—including CNN_C1u and CNN_C23u— trained on DisProt sequences (dominated by long sites) exhibit a clear preference for short sites in the DBsh dataset. Furthermore, CNN_TR08u, trained against very short sites (≤25 residues), yielded comparable performance on short and long sites in DBsh, and CNN_DBs, trained against the DBs dataset dominated by long sites, yielded much higher AUC values in identifying short sites in the CAID datasets. Thus, while we cannot rule out the binding site size as a minor factor influencing the performance of these predictors, we can conclude that it is not the main factor.

**Table 3.** The AUC results in identifying short (<=70 AA) and long (>70 AA) binding sites for our 11 predictors, plus the 4 CNN predictors against the three test sets, CAID1uh, CAID2&3uh, and DBsh.

|  |  | CAID1uh |  | CAID23uh |  | DBsh |  |
| --- | --- | --- | --- | --- | --- | --- | --- |
|  | Group | Short | Long | Short | Long | Short | Long |
| AlphaFold-binding | A |  |  | 0.800 | 0.785 | 0.642 | 0.372 |
| ANCHOR-2 |  | 0.650 | 0.762 | 0.678 | 0.766 | 0.496 | 0.369 |
| CNN_C1u | A |  |  | 0.757 | 0.822 | 0.671 | 0.518 |
| CNN_C23u | A | 0.743 | 0.793 |  |  | 0.702 | 0.522 |
| CNN_DBs | B | 0.693 | 0.523 | 0.692 | 0.529 |  |  |
| CNN_TR2008u | B | 0.748 | 0.665 | 0.772 | 0.691 | 0.779 | 0.743 |
| DeepDISObind-protein | A |  |  | 0.722 | 0.763 | 0.544 | 0.366 |
| DeepDRPBind-protein | A |  |  | 0.683 | 0.729 | 0.646 | 0.564 |
| DisoRDPbind-protein | A | 0.659 | 0.762 | 0.693 | 0.776 | 0.483 | 0.350 |
| DRPBind-protein | A |  |  | 0.665 | 0.754 | 0.467 | 0.322 |
| fMoRFpred | B | 0.582 | 0.533 | 0.592 | 0.552 | 0.615 | 0.535 |
| OPAL | B | 0.772 | 0.629 | 0.769 | 0.701 | 0.788 | 0.722 |
| MoRFchibi | B | 0.612 | 0.521 | 0.629 | 0.549 | 0.624 | 0.610 |
| MoRFchibi-light | C | 0.738 | 0.699 | 0.776 | 0.774 | 0.721 | 0.587 |
| MoRFchibi-web | C | 0.732 | 0.665 | 0.784 | 0.759 | 0.728 | 0.619 |

### 6. Investigating data features

Each class (binding vs. non-binding) can have features specific to its sequences. In addition, some tools (e.g., MoRFchibi and ANCHOR-2) rely solely on non-sequence features averaged over a window of a given size. In this section, we examine the consistency of conservation and disorder levels between the training and test datasets and assess their potential contribution to differences in predictor performance. More specifically, a feature is considered to contribute to such performance discrepancies if the following pattern holds: test datasets that yield high AUC values exhibit feature levels similar to those in the training dataset, whereas test datasets that produce low AUC values show dissimilar feature levels — consistent with the observations reported in Sections 3 and 4:

A. In Class 0, the feature levels are broadly similar across all datasets (Section 4).
B. In Class 1:
  1. Feature levels are comparable between the CAID1u(h) and CAID2&3u(h) datasets (Section 4).
  2. Feature levels differ clearly between TR2008u and the two CAID datasets (Section 3).
  3. Feature levels differ clearly between TR2008u and the DBs(h) dataset (Section 3).
  4. Feature levels show the greatest divergence between the DBs(h) dataset and the two CAID datasets (Section 3).

### 6.1 Conservation

As functional elements, the sequences of IDR-protein binding sites are under selective pressure and are more important than those of other disordered regions (e.g., Linkers). Supplementary Figure 2-A presents violin plots of LIST-S2 [67] conservation scores for IDR-protein binding sites (class 1) in the CAID1u, CAID2&3u, DBs, and TR2008u datasets, as well as for non-IDR-protein binding residues (class 0), compared with the conservation of the CAIDs IDR and folded (PDB) regions. The results are inconsistent with observation (B.1), as we observe no conservation similarity among IDR-protein binding sites across the CAID datasets. Consequently, we can rule out conservation as a major factor influencing the performance of these predictors.

### 6.2 Disorderness

It was shown that MoRFs, a subcategory of IDR protein-binding sites, exhibit reduced disorder propensity (disorderness) compared to their surrounding IDR regions [15]. **Supplementary Figure 2-B** plots IUPred-3 disorderness score violin plots for IDR-protein binding sites (class 1) in the CAID1u, CAID2&3u, DBs, and TR2008 datasets, as well as those for non-IDR-protein binding residues (class 0) compared to the disorderness of the CAIDs IDR and folded (PDB) regions. Results show that the disorderness of class 0 is about the same, in line with observation (A). For class 1, similarity between the CAID1u(h) and the CAID2&3u(h) datasets, observation (B.1), a clear difference between TR2008u and the CAID datasets (B.2), a clear difference between the DBsh and the TR2008u (B.3), and the sharpest difference between DBsh and the CAID datasets (B.4). Consequently, we can conclude that disorderness is the main feature that influences IDR-protein binding computational prediction tools.

This conclusion can also explain other unanticipated results: First, the limited overlap between the disorderness on the DBs class 1 residues and the CAID class 1 residues can explain the abnormal shape of the ROC curves for predictors trained using DisProt sequences observed in **section 2, Figure 1**, as the disorderness of many DBs class 1 residues is more consistent with that of the class 0 residues in the DisProt sequences, **Supplementary Figure 2**. Second, when we compare the disorderness for short (<=70 AA) and long (>70 AA) in our three test datasets, CAID1uh, CAID2&3uh, and DBsh, **Table 4** and **Figure 4**, we can see that the short binding sites in the DBsh dataset are more disordered than those that are long. Consequently, tools trained on datasets sourced from the DisProt database, which contain more disordered class 1 residues, achieve higher AUCs for identifying more disordered short sites, thereby explaining the observation in **section 5**. Finally, CNN_DBs trained on binding sites with lower disorderness values are better able to identify the less-disordered short binding sites in the CAID datasets, as shown in **section 5**.

**Table 4.** The mean values of the IUPred3 disorderness for class 0 and class 1 in the three training datasets and the three test (with suffix h) datasets (all, short, and long). The mean IUPred scores for the TR2008 class 0 is 0.332, class 1 is 0.377, IDR is 0.495, and folded domains (PDB) is 0.200

|  | CAID1u-1 | CAID2&3u-1 | Dbs-1 | CAID1u-0 | CAID2&3u-0 | Dbs-0 |
| --- | --- | --- | --- | --- | --- | --- |
| **\*-all** | 0.518 | 0.508 | 0.258 | 0.324 | 0.320 | 0.356 |
| **\*h-all** | 0.516 | 0.509 | 0.258 | 0.324 | 0.324 | 0.356 |
| **\*h-short** | 0.487 | 0.448 | 0.334 | 0.324 | 0.324 | 0.356 |
| **\*h-long** | 0.525 | 0.521 | 0.244 | 0.324 | 0.324 | 0.356 |

**Figure 4.**
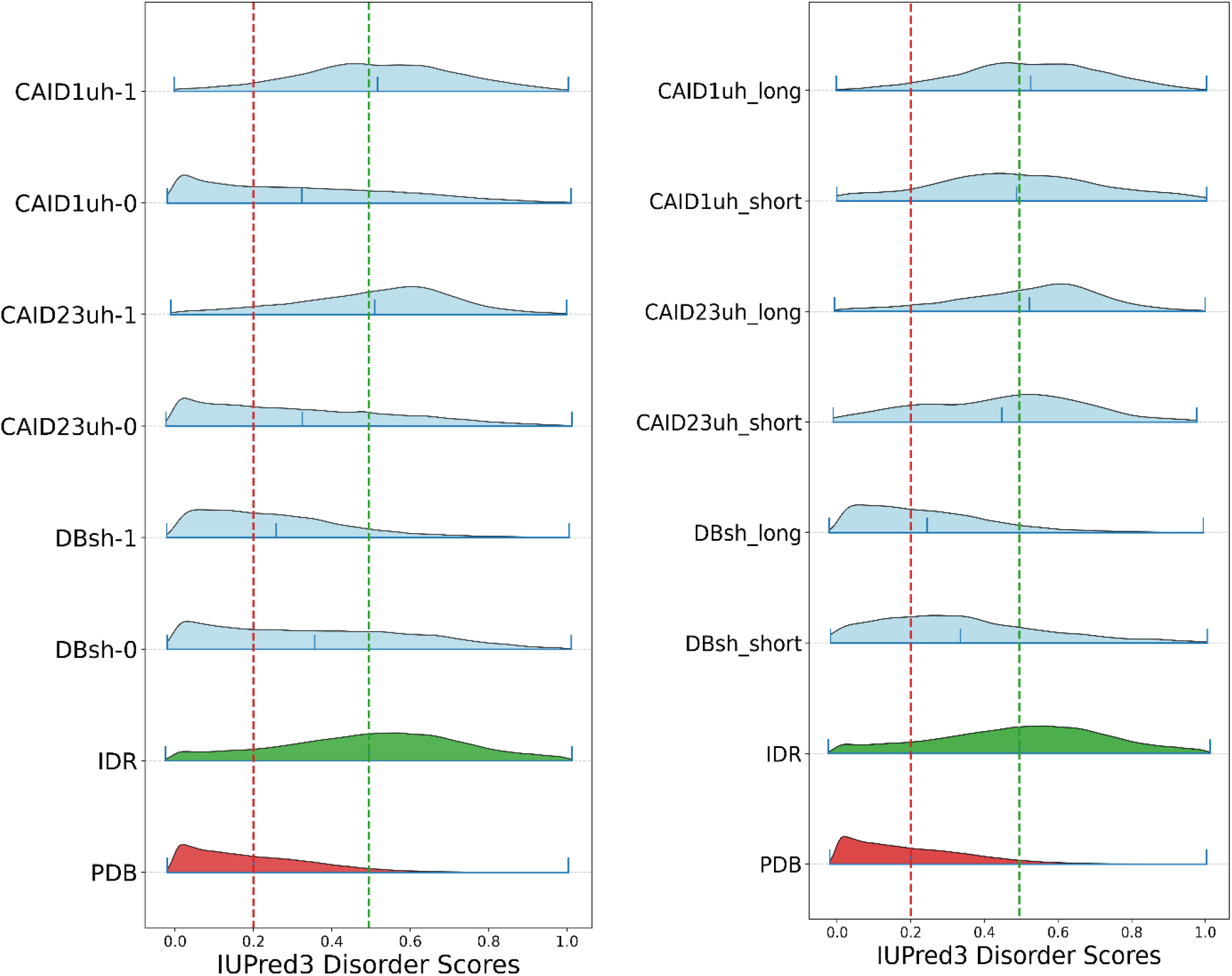
IUPred3 disorderness violin plots for the three test datasets class 0 and class 1, (all, short, and long) binding sites compared to those of the IDR and folded domains (PDB) in the CAID datasets.

This leaves us with the ANCHOR-2 predictor, optimized against very short IDR-protein binding sites sourced from PDB structures (i.e., less disordered), and it performs in line with Group A predictors trained against highly disordered binding sites in the DisProt database sourced from the literature. ANCHOR-2 is a “simple biophysics-based model” [35] with only four optimizable parameters learned from the TR2017 training dataset. Its final score for a residue at position k is the Sigmoid of the product of two sub-scores. First, the energy gain for the residue binding with a predefined generic globular domain, and the second value is the average IUPred [35] disorderness score. Thus, the unexpected performance of ANCHOR-2 can be explained by its second sub-score: higher disorderness directly increases this sub-score and, in turn, the final ANCHOR-2 score. Accordingly, ANCHOR-2 has been constantly performing in line with predictors in Group A that are trained against highly disordered class 1 residues, not Group B, further supporting our conclusion that disorderness is the main feature influencing the prediction of IDR-protein binding sites, and thus resulting in the inconsistencies in predictors’ performance against the test dataset with different class 1 disorderness levels.

## 7. Discussion

This study identifies a fundamental, previously overlooked “hidden disorder divide” that causes systematic performance inconsistencies among IDR protein-binding computational tools across major high-quality, manually curated datasets. Beyond methodological flaws or variations in machine-learning architectures, these discrepancies are significantly influenced by differences in the intrinsic disorder propensity of the binding sites. Our analysis of 11 established predictors reveals a clear split in performance across training datasets. Predictors trained on DisProt sequences, such as DeepDISObind-protein and AlphaFold-binding (Group A), excel on literature-based CAID benchmarks but suffer a dramatic performance drop (average ~31 AUC points) when tested against PDB-derived datasets (DBs). In contrast, predictors trained on older, PDB-derived data, including MoRFchibi and OPAL, show a modest preference for the DBs over the two CAID datasets.

Our experiments with four control CNN predictors show that the observed effects are not solely due to the model architecture but rather reflect properties of the datasets themselves. Class-specific residue merging further confirmed that differences in binding residues are the primary drivers of performance gaps, whereas non-binding residues have a negligible impact on performance differences. The root of this “divide” is a distinct selection bias in how binding sites are curated. Literature-based annotations (DisProt) preferentially capture highly disordered binding sites (mean IUPred3 score ~0.51), characterized by various experimental techniques. PDB-based annotations predominantly capture less-disordered binding sites (mean scores ~0.26 to ~0.38) that are amenable to crystallization in complexes with partners. Other factors, such as binding site length and sequence conservation, were found to play a minor, if any, role in explaining these inconsistencies.

Interestingly, ANCHOR-2 performs more like DisProt-trained tools despite its PDB-based training. This is explained by its biophysics-based model, which explicitly incorporates disorder scores, such that higher disorder propensity directly increases its prediction confidence.

### 8. Conclusions and Future Directions

The practical implications of these findings are significant for the development and evaluation of future predictors. They underscore the urgent need for more balanced reference datasets that better capture the full spectrum of disorder propensity in IDR-mediated binding. Future models should explicitly account for disorder heterogeneity, for example, through disorder-balanced training sets or stratified sampling strategies. To ensure a complete and unbiased assessment of a tool’s utility, benchmarks should report performance separately across different disorder regimes, rather than relying on a single, potentially skewed dataset.

Ultimately, the “hidden disorder divide” mirrors the genuine biological diversity in how disordered regions interact with their cellular partners. Recognizing this divide is essential for deepening our understanding of IDR functionality and for developing next-generation predictors that can accurately identify binding sites across the entire disorder spectrum. Finally, diagnostic techniques such as the control-model and class-specific residue merging approaches introduced here could prove valuable more broadly, especially when test sets originate from heterogeneous sources. These practices should become standard in the field.

