## Supplemental file for "The Hidden Disorder Divide: Reconciling Benchmark Inconsistencies in Intrinsically Disordered Protein Binding Site Prediction"

Supplement Note 1

**Datasets**

**TR2008:** The TR2008 training dataset was constructed before the emergence of manually curated IDR–protein binding databases. Its authors selected PDB structures that capture biologically relevant interactions between a large, structured protein domain (>70 residues) and a short peptide (≤25 residues). These peptides were classified as MoRFs if in silico predictors indicated they were disordered in isolation. The resulting MoRF regions were then mapped to UniProtKB sequences, with the restriction that only one MoRF was annotated per protein sequence. TR2008 has 245,984 residues in 421 sequences from 389 unique UniProt IDs. Class 1, 421 binding sites, includes 5,396 residues; the remaining 240,588 residues are class 0 (for more details, see [1]).

This one-MoRF-per-sequence constraint introduced annotation conflicts and redundancies, particularly among highly similar sequences that received inconsistent labels [2]. For example, two 212-residue Hepatitis B virus capsid proteins, UniProt entries Q461C5 and Q68032, share 96.7% sequence identity. Nevertheless, their MoRF annotations differ: residues 47–56 are marked as a MoRF in Q461C5 but not in Q68032, whereas residues 116–124 are annotated as a MoRF in Q68032 but not in Q461C5.

```
########################################
# Program: needle
#    -gapopen 10.0
#    -gapextend 0.5
#    -endopen 10.0
#    -endextend 0.5
#    -aformat3 pair
#    -sprotein1
#    -sprotein2
# Align_format: pair
# Report_file: stdout
########################################

Q461C5             1 MQLFDLCLIISCSCPTVQASKLCLGWLWGMDIDPYKEFGATVELLSFLPS     50
                     ||||.|||||||||||||||||||||||||||||||||||||||||||||
Q68032             1 MQLFHLCLIISCSCPTVQASKLCLGWLWGMDIDPYKEFGATVELLSFLPS     50

Q461C5            51 DFFPSVRDLLDTASALYRDALESPEHCSPHHTALRQAILCWGELMTLATW    100
                     ||||||||||||||||||:|||||||||||||||||||||||:|||||||
Q68032            51 DFFPSVRDLLDTASALYREALESPEHCSPHHTALRQAILCWGDLMTLATW    100

Q461C5           101 VGVNLEDPASRDLVVSYVNTNMGLKFRQLLWFHISCLTFGRETVIEYLVS    150
                     ||||||||||||||||||||||||||:|||||||||||||||||||||||
Q68032           101 VGVNLEDPASRDLVVSYVNTNMGLKFKQLLWFHISCLTFGRETVIEYLVS    150

Q461C5           151 FGVWIRTPPAYRPPNAPILSTLPENTVVRRRGRSPRRRTPSPRRRRSQSP    200
                     ||||||||||||||||||||||||.|||||||||||||||||||||||||
Q68032           151 FGVWIRTPPAYRPPNAPILSTLPETTVVRRRGRSPRRRTPSPRRRRSQSP    200

Q461C5           201 RRRRSQSPASQC    212
                     |||||||..|||
Q68032           201 RRRRSQSRESQC    212
#---------------------------------------
#---------------------------------------
```

**TS2008:** It was constructed jointly with TR2008 and then separated from it with <30% identity. It includes 419 sequences totaling 258829 residues. It includes 419 binding sites, each ≤25 residues, totalling 5,153.

**TS2012:** collected by the authors of MoRFpred, using an approach similar to that used by TR2008 and TS2008, but from PDB structures deposited between 2008 and 2021. It is limited to 45 protein sequences totaling 37,533 residues, including 626 residues across 45 binding sites.

**TR2017:** Class 1, obtained from the DIBS database in 2017 [3], comprises 250 short linear binding sites (<30 residues; longer sites were excluded), totaling 3,858 residues. Class 0 consists of 663,433 residues across 10,914 protein regions split into two subsets:

1. 3,033 randomly selected monomeric single-domain PDB structures with <20% flexible residues (533,342 residues), and
2. 10,152 randomly chosen intracellular human protein regions lacking transmembrane segments or Pfam domains (156,744 residues), with length distributions matched to the positive DIBS segments (for details, see [4]).

**Supplement Note 2**

**CNN prediction model**

Each CNN predictor comprises three 1D convolutional layers (with two 1D average-pooling layers inserted after the first two convolutions) followed by two fully connected layers. The architecture is as follows:

1. 1D convolutional layer (input length 510, kernel size CKZ, K filters) → ReLU → average pooling (kernel size APKZ)
2. 1D convolutional layer (kernel size CKZ, K + D_inc filters) → ReLU → average pooling (kernel size APKZ)
3. 1D convolutional layer (kernel size CKZ, K + 2·D_inc filters) → ReLU
4. Fully connected layer (FC1, output size FC1Z) → ReLU
5. Fully connected layer (FC2, output size 10)

Input sequences are symmetrically padded on each side with 250 “blank” residues (all feature values = 0). Prediction proceeds in sliding windows of 510 residues: the first window covers actual residue positions p = 1 to p = 10, with residues p-250 to p+259 supplied as input (using the K = 7 features per residue). The 10 outputs from FC2 correspond to residues p through p+9. The window is then shifted by 10 residues (p ← p + 10), and the process repeats until the entire sequence is covered.

The seven input features, sourced from IPA, are:

- Two affinity features are derived from P, a 20 × 20 energy predictor matrix [5]. For each position p between 1 and the last in the sequence, one of the affinity features is the sum of the energy in P between the residue at p and the w1=10 residues on each side of p. The second is the sum of this energy between w1 and w2=w1+25, from p + w1 to p + w2 on each side of p.
- The F5 features are the frequency of each of the 20 standard amino acids in each of the five classes in the DisProt/CAID datasets: IDR, nucleic binding, protein binding, and ‘Linker’ in the DisProt data and ‘PDB’ annotation in the combined CAID1 and CAID2 sequences, divided by the frequency of that amino acid in the entire dataset

**Supplement Table S1**

**Dataset Compositions:** The number of sequences, the number of binding sites, total residue counts, and per-class counts for each dataset.

| Dataset | Total Seq. | Total Residues | Seq. with sites | Number of Sites | Class 0 (non-binding) | Class 1 (binding) | Masked residues |
| --- | --- | --- | --- | --- | --- | --- | --- |
| CAID1u | 599 | 315,318 | 208 | 234 | 91,055 | 19,656 | 112 |
| CAID1uh | 599 | 315,318 | 201 | 225 | 87,882 | 19,213 | 217 |
| CAID23u | 612 | 403,677 | 86 | 93 | 53,703 | 9,409 | 441 |
| CAID23uh | 612 | 403,677 | 83 | 89 | 45,945 | 9,246 | 1,274 |
| DBs | 757 | 341,749 | 757 | 797 | 263,001 | 78,748 | 0 |
| DBsh | 757 | 341,749 | 757 | 809 | 262,658 | 78,608 | 483 |
| TR2008u | 308 | 170,121 | 308 | 308 | 165,871 | 3,978 | 272 |
| VDS | 427 | 254,314 | 427 | 472 | 226,099 | 28,215 | 0 |
| VA_1 | 197 | 125,755 | 197 | 242 | 100,395 | 25,360 | 0 |
| VA_2 | 230 | 128,559 | 230 | 230 | 125,704 | 2,855 | 0 |

**Supplementary Table S2**

AUC values for identifying IDR-protein binding sites across all 11 predictors and the four CNN models, evaluated on our three test datasets processed by the HAM software to mask local homologies. Tools constructed after the CAID round 1 event are not evaluated on the CAID1uh dataset.

|  | **Predictor** | **Group** | **AUC** | | |
| --- | --- | --- | --- | --- | --- |
|  |  |  | **CAID1uh** | **CAID2&3uh** | **DBsh** |
| 1 | AlphaFold-binding | A |  | 0.788 | 0.412 |
| 2 | ANCHOR-2 |  | 0.737 | 0.752 | 0.389 |
|  | CNN_C1u | A |  | 0.811 | 0.541 |
|  | CNN_C23u | A | 0.782 |  | 0.549 |
|  | CNN_DBs | B | 0.562 | 0.556 |  |
|  | CNN_TR08u | B | 0.684 | 0.704 | 0.748 |
| 3 | DeepDISObind-protein | A |  | 0.756 | 0.393 |
| 4 | DeepDRPBind-protein | A |  | 0.721 | 0.577 |
| 5 | DisoRDPbind-protein | A | 0.739 | 0.762 | 0.370 |
| 6 | DRPBind-protein | A |  | 0.739 | 0.344 |
| 7 | fMoRFpred | B | 0.544 | 0.558 | 0.547 |
| 8 | OPAL | B | 0.661 | 0.712 | 0.732 |
| 9 | MoRFchibi | B | 0.542 | 0.562 | 0.612 |
| 10 | MoRFchibi-light | C | 0.708 | 0.774 | 0.607 |
| 11 | MoRFchibi-web | C | 0.680 | 0.763 | 0.635 |

**Supplementary Table S3**

**Table S3 (A):** AUC values from the CNN_C1u model against the CAID2&3uh and DBsh test datasets for the 21 instances ranked 20th to 40th highest AUCs on the validation set.

|  | **AUCs** | | |
| --- | --- | --- | --- |
| **Rank** | **CAID1uh** | **CAID23uh** | **DBsh** |
| 20 |  | 0.820 | 0.554 |
| 21 |  | 0.821 | 0.567 |
| 22 |  | 0.815 | 0.530 |
| 23 |  | 0.817 | 0.557 |
| 24 |  | 0.822 | 0.515 |
| 25 |  | 0.802 | 0.510 |
| 26 |  | 0.825 | 0.522 |
| 27 |  | 0.816 | 0.569 |
| 28 |  | 0.819 | 0.529 |
| 29 |  | 0.825 | 0.552 |
| **30** |  | 0.811 | 0.541 |
| 31 |  | 0.821 | 0.510 |
| 32 |  | 0.824 | 0.539 |
| 33 |  | 0.816 | 0.523 |
| 34 |  | 0.827 | 0.538 |
| 35 |  | 0.816 | 0.534 |
| 36 |  | 0.817 | 0.523 |
| 37 |  | 0.815 | 0.558 |
| 38 |  | 0.818 | 0.532 |
| 39 |  | 0.818 | 0.513 |
| 40 |  | 0.816 | 0.517 |

**Table S3 (B):** AUC values from the CNN_C23u model against the CAID1uh and DBsh test datasets for the 21 instances ranked 20th to 40th highest AUCs on the validation set.

|  | **AUCs** | | |
| --- | --- | --- | --- |
| **Rank** | **CAID1uh** | **CAID2&3uh** | **DBsh** |
| 20 | 0.773 |  | 0.564 |
| 21 | 0.774 |  | 0.574 |
| 22 | 0.768 |  | 0.560 |
| 23 | 0.778 |  | 0.546 |
| 24 | 0.786 |  | 0.565 |
| 25 | 0.773 |  | 0.573 |
| 26 | 0.770 |  | 0.583 |
| 27 | 0.781 |  | 0.570 |
| 28 | 0.767 |  | 0.578 |
| 29 | 0.756 |  | 0.590 |
| **30** | **0.782** |  | **0.550** |
| 31 | 0.773 |  | 0.575 |
| 32 | 0.763 |  | 0.619 |
| 33 | 0.774 |  | 0.578 |
| 34 | 0.781 |  | 0.589 |
| 35 | 0.773 |  | 0.560 |
| 36 | 0.775 |  | 0.579 |
| 37 | 0.768 |  | 0.593 |
| 38 | 0.776 |  | 0.571 |
| 39 | 0.768 |  | 0.568 |
| 40 | 0.774 |  | 0.565 |

**Table S3 (C):** AUC values from the CNN_DBs model against the CAID1uh and CAID2&3uh test datasets for the 21 instances ranked 20th to 40th highest AUCs on the validation set.

|  | **AUCs** | | |
| --- | --- | --- | --- |
| **Rank** | **CAID1uh** | **CAID2&3uh** | **DBsh** |
| 20 | 0.575 | 0.562 |  |
| 21 | 0.576 | 0.564 |  |
| 22 | 0.562 | 0.541 |  |
| 23 | 0.554 | 0.548 |  |
| 24 | 0.563 | 0.549 |  |
| 25 | 0.556 | 0.538 |  |
| 26 | 0.553 | 0.532 |  |
| 27 | 0.554 | 0.530 |  |
| 28 | 0.567 | 0.551 |  |
| 29 | 0.573 | 0.552 |  |
| **30** | 0.562 | 0.556 |  |
| 31 | 0.563 | 0.545 |  |
| 32 | 0.575 | 0.550 |  |
| 33 | 0.550 | 0.527 |  |
| 34 | 0.548 | 0.527 |  |
| 35 | 0.553 | 0.542 |  |
| 36 | 0.551 | 0.525 |  |
| 37 | 0.542 | 0.521 |  |
| 38 | 0.549 | 0.517 |  |
| 39 | 0.560 | 0.533 |  |
| 40 | 0.542 | 0.517 |  |

**Table S3 (D):** AUC values from the CNN_TR08u model against the CAID1uh, CAID2&3uh, and DBsh test datasets for the 21 instances ranked 20th to 40th highest AUCs on the validation set.

| **Rank** | **CAID1uh** | **CAID2&3uh** | **DBsh** |
| --- | --- | --- | --- |
| 20 | 0.676 | 0.700 | 0.776 |
| 21 | 0.696 | 0.716 | 0.767 |
| 22 | 0.687 | 0.698 | 0.737 |
| 23 | 0.685 | 0.707 | 0.772 |
| 24 | 0.684 | 0.696 | 0.757 |
| 25 | 0.683 | 0.718 | 0.758 |
| 26 | 0.693 | 0.710 | 0.753 |
| 27 | 0.685 | 0.716 | 0.746 |
| 28 | 0.683 | 0.700 | 0.764 |
| 29 | 0.696 | 0.712 | 0.723 |
| **30** | 0.684 | 0.704 | 0.748 |
| 31 | 0.665 | 0.684 | 0.769 |
| 32 | 0.669 | 0.700 | 0.780 |
| 33 | 0.684 | 0.708 | 0.782 |
| 34 | 0.689 | 0.711 | 0.740 |
| 35 | 0.652 | 0.677 | 0.783 |
| 36 | 0.684 | 0.694 | 0.725 |
| 37 | 0.678 | 0.704 | 0.775 |
| 38 | 0.680 | 0.684 | 0.762 |
| 39 | 0.659 | 0.672 | 0.790 |
| 40 | 0.669 | 0.687 | 0.772 |

**Supplementary Table S4**

**Supplementary Table S4:** Predictors' performance in separating protein-binding sites, class 1, in the CAID2&3uh and the DBsh datasets from a class 0 that is limited to disordered non-binding IDRs (IDR) or folded protein regions (PDB) in the CAID2&3uh dataset.

|  | **CAID2&3uh** | | **DBsh** | |
| --- | --- | --- | --- | --- |
|  | **PDB** | **IDR** | **PDB** | **IDR** |
| **AlphaFold-binding** | 0.961 | 0.556 | 0.755 | 0.220 |
| **ANCHOR-2** | 0.886 | 0.560 | 0.584 | 0.222 |
| **CNN_C1u** | 0.938 | 0.585 | 0.771 | 0.315 |
| **CNN_TR2008u** | 0.737 | 0.496 | 0.780 | 0.573 |
| **DeepDISObind-protein** | 0.915 | 0.539 | 0.657 | 0.204 |
| **DeepDRPBind-protein** | 0.794 | 0.524 | 0.699 | 0.404 |
| **DisoRDPbind-protein** | 0.908 | 0.570 | 0.600 | 0.211 |
| **DRPBind-protein** | 0.891 | 0.533 | 0.572 | 0.175 |
| **fMoRFpred** | 0.576 | 0.529 | 0.569 | 0.523 |
| **OPAL** | 0.731 | 0.531 | 0.766 | 0.590 |
| **MoRFchibi** | 0.539 | 0.526 | 0.572 | 0.560 |
| **MoRFchibi-light** | 0.851 | 0.578 | 0.714 | 0.430 |
| **MoRFchibi-web** | 0.817 | 0.598 | 0.699 | 0.474 |

**Supplementary Table S5:** the AUC and Δ AUC values for the eleven predictors, CNN_C1u, and CNN_TR08u against the CAID2&3uh and the DBsh test datasets in two scenarios: (A) Targeting class 1 by merging class 0, and (B) targeting class 0 by merging class 1.

| **Class 0 merged** | **Group** | **CAID2&3uh** | **DBsh** | Δ AUC |
| --- | --- | --- | --- | --- |
| AlphaFold-binding | A | 0.759 | 0.467 | 0.292 |
| ANCHOR-2 |  | 0.735 | 0.416 | 0.319 |
| CNN_C1u | A | 0.799 | 0.573 | 0.227 |
| CNN_TR08u | B | 0.702 | 0.751 | -0.049 |
| DeepDISObind-protein | A | 0.740 | 0.436 | 0.305 |
| DeepDRPBind-protein | A | 0.707 | 0.601 | 0.106 |
| DisoRDPbind-protein | A | 0.746 | 0.405 | 0.340 |
| DRPBind-protein | A | 0.724 | 0.381 | 0.342 |
| fMoRFpred | B | 0.556 | 0.550 | 0.007 |
| OPAL | B | 0.704 | 0.743 | -0.038 |
| MoRFchibi | B | 0.569 | 0.602 | -0.032 |
| MoRFchibi-light | C | 0.769 | 0.621 | 0.148 |
| MoRFchibi-web | C | 0.762 | 0.639 | 0.123 |
| **Class 1 merged** | **Group** | **CAID2&3uh** | **DBsh** | Δ AUC |
| AlphaFold-binding | A | 0.532 | 0.442 | 0.090 |
| ANCHOR-2 |  | 0.469 | 0.423 | 0.046 |
| CNN_C1u | A | 0.617 | 0.566 | 0.051 |
| CNN_TR08u | B | 0.747 | 0.743 | 0.004 |
| DeepDISObind-protein | A | 0.496 | 0.428 | 0.068 |
| DeepDRPBind-protein | A | 0.629 | 0.588 | 0.041 |
| DisoRDPbind-protein | A | 0.465 | 0.407 | 0.058 |
| DRPBind-protein | A | 0.441 | 0.382 | 0.059 |
| fMoRFpred | B | 0.552 | 0.548 | 0.005 |
| OPAL | B | 0.746 | 0.728 | 0.017 |
| MoRFchibi | B | 0.591 | 0.608 | -0.018 |
| MoRFchibi-light | C | 0.645 | 0.624 | 0.022 |
| MoRFchibi-web | C | 0.654 | 0.648 | 0.006 |

**Supplementary Table S6**: the AUC and Δ AUC values for the seven predictors, CNN_C23u, and CNN_TR08u against the CAID1uh and the DBsh test datasets in two scenarios: (A) Targeting class 1 by merging class 0, and (B) targeting class 0 by merging class 1.

| **Class 0 merged** | **Group** | **CAID1uh** | **DBsh** | Δ AUC |
| --- | --- | --- | --- | --- |
| ANCHOR-2 |  | 0.717 | 0.414 | 0.304 |
| CNN_C23u | A | 0.771 | 0.569 | 0.202 |
| CNN_TR08u | B | 0.695 | 0.736 | -0.041 |
| DisoRDPbind-protein | A | 0.716 | 0.400 | 0.316 |
| fMoRFpred | B | 0.544 | 0.547 | -0.003 |
| OPAL | B | 0.670 | 0.723 | -0.052 |
| MoRFchibi | B | 0.550 | 0.603 | -0.053 |
| MoRFchibi-light | C | 0.707 | 0.612 | 0.095 |
| MoRFchibi-web | C | 0.681 | 0.636 | 0.045 |
| **Class 1 merged** | **Group** | **CAID1uh** | **DBsh** | Δ AUC |
| ANCHOR-2 |  | 0.496 | 0.449 | 0.047 |
| CNN_C23u | A | 0.625 | 0.591 | 0.035 |
| CNN_TR2008u | B | 0.717 | 0.740 | -0.023 |
| DisoRDPbind-protein | A | 0.489 | 0.433 | 0.055 |
| fMoRFpred | B | 0.546 | 0.546 | 0.000 |
| OPAL | B | 0.703 | 0.722 | -0.019 |
| MoRFchibi | B | 0.584 | 0.601 | -0.017 |
| MoRFchibi-light | C | 0.634 | 0.627 | 0.007 |
| MoRFchibi-web | C | 0.645 | 0.644 | 0.001 |

**Supplementary Table S7**: the AUC and Δ AUC values for the eleven predictors, CNN_DBs, and CNN_TR08u against the CAID1uh and the CAID2&3uh test datasets in two scenarios: (A) Targeting class 1 by merging class 0, and (B) targeting class 0 by merging class 1.

| **Class 0 merged** | **Group** | **CAID1uh** | **CAID2&3uh** | **Δ AUC** |
| --- | --- | --- | --- | --- |
| ANCHOR-2 |  | 0.738 | 0.752 | -0.014 |
| CNN_DBs | B | 0.580 | 0.540 | 0.040 |
| CNN_TR2008u | B | 0.700 | 0.691 | 0.009 |
| DisoRDPbind-protein | A | 0.737 | 0.764 | -0.026 |
| fMoRFpred | B | 0.546 | 0.556 | -0.010 |
| OPAL | B | 0.682 | 0.695 | -0.012 |
| MoRFchibi | B | 0.541 | 0.562 | -0.021 |
| MoRFchibi-light | C | 0.716 | 0.767 | -0.051 |
| MoRFchibi-web | C | 0.684 | 0.760 | -0.076 |
| **Class 1 merged** | **Group** | **CAID1uh** | **CAID2&3uh** | Δ AUC |
| ANCHOR-2 |  | 0.742 | 0.742 | -0.001 |
| CNN_DBs | B | 0.548 | 0.582 | -0.035 |
| CNN_TR2008u | B | 0.681 | 0.710 | -0.029 |
| DisoRDPbind-protein | A | 0.748 | 0.745 | 0.003 |
| fMoRFpred | B | 0.547 | 0.551 | -0.004 |
| OPAL | B | 0.664 | 0.703 | -0.038 |
| MoRFchibi | B | 0.548 | 0.548 | 0.001 |
| MoRFchibi-light | C | 0.724 | 0.739 | -0.015 |
| MoRFchibi-web | C | 0.705 | 0.711 | -0.007 |

**Supplementary Figure 1**

| 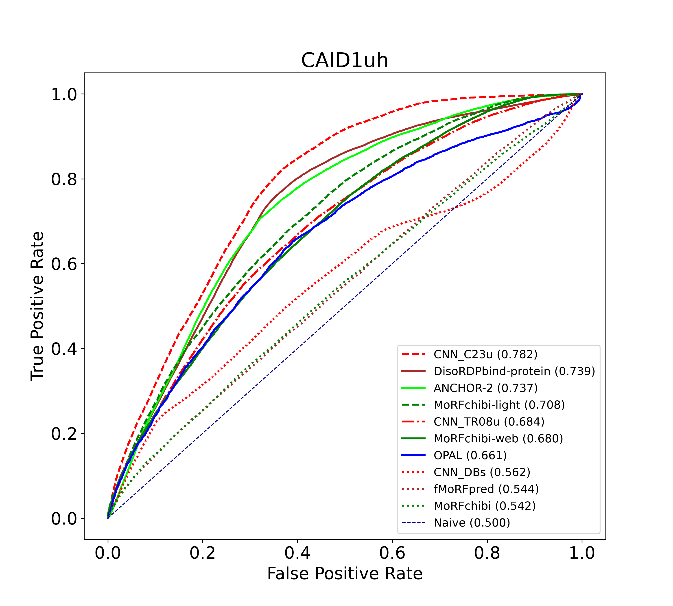 | 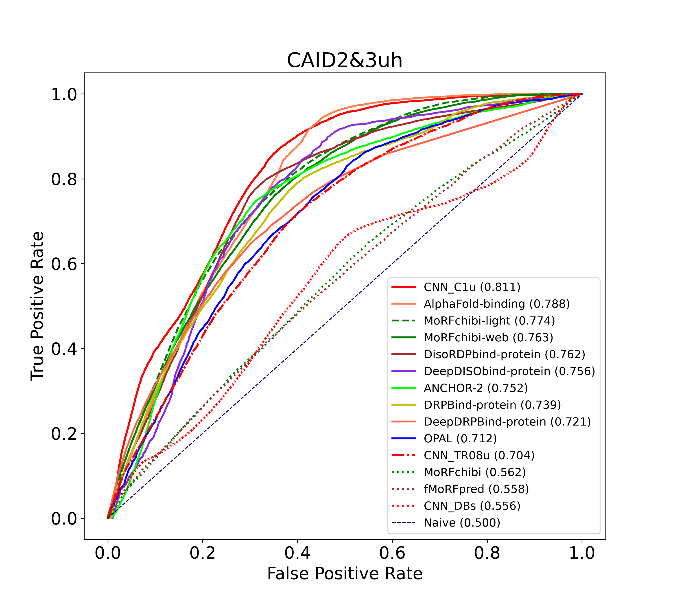 |
| --- | --- |
| 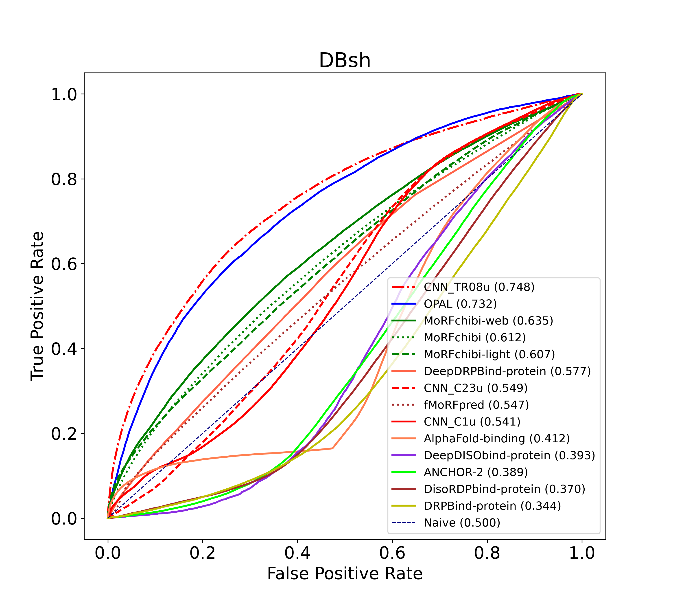 | 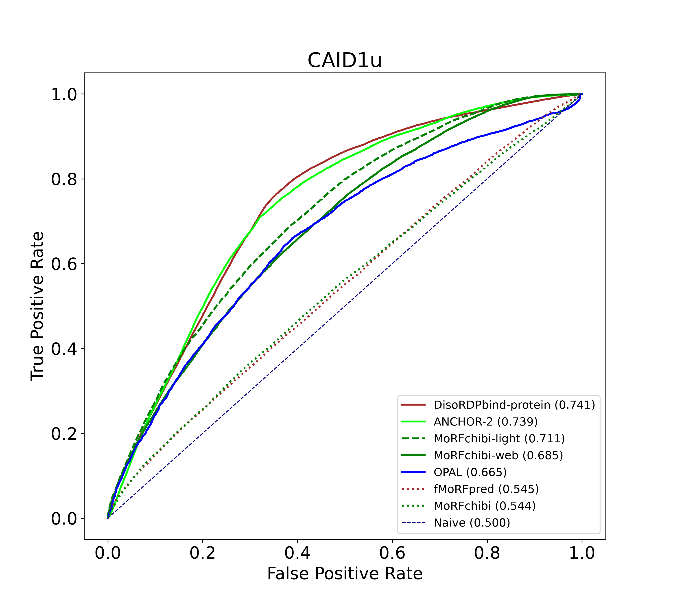 |
| **Figure S1:** (a, b, and c) are the ROC curves for the eleven IDR-protein binding predictors and four CNN_* new predictors against the CAID1uh, CAID2&3uh, and DBsh datasets. (d) is the ROC curve for the seven predictors that participated in the CAID round 1, evaluated on the CAID1u test dataset. | |

**Supplementary Figure 2**

| 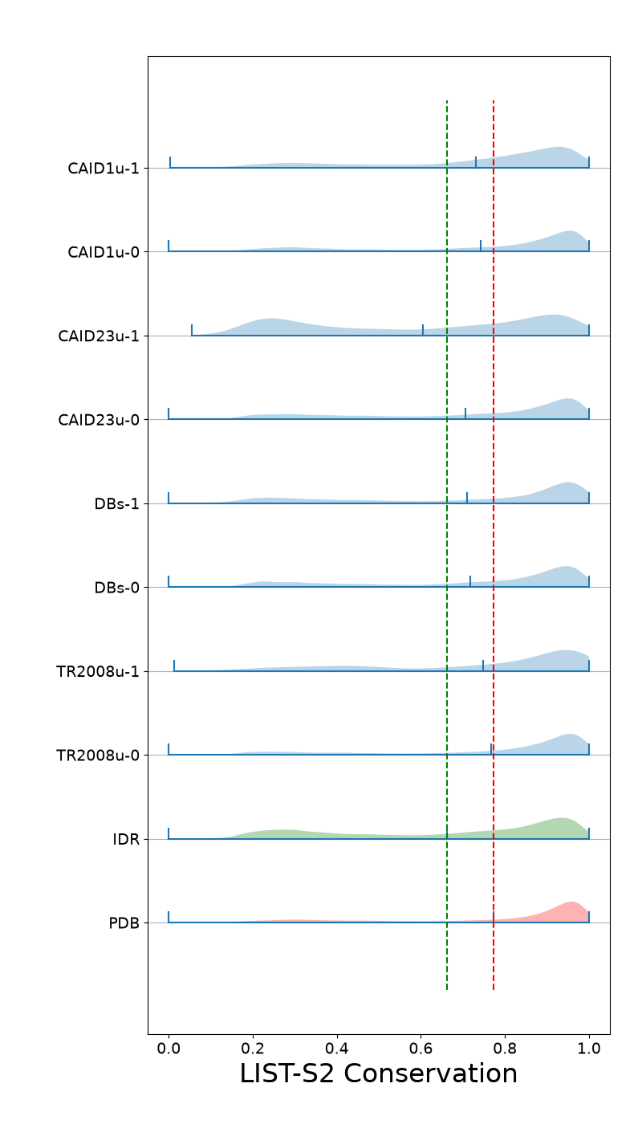 | 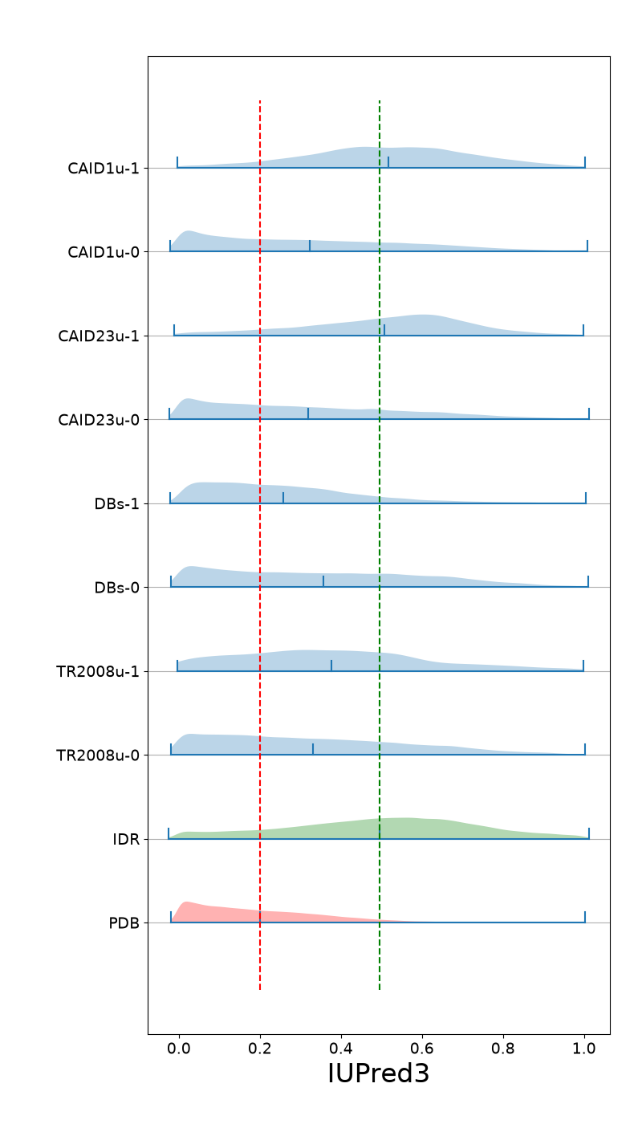 |
| --- | --- |
| **Supplementary Figure 2:** Violin plots for the LIST-S2 conservation (left) and IUPred-3 short disorderness (right) score for the IDR-protein binding sites, class 1, and the remaining residues, class 0 in CAID1u, CAID2&3u, DBs, and TR2008u, and IDR and PDB regions in the combined CAID datasets. | |
